# Comprehensive structural analysis reveals broad-spectrum neutralizing antibodies against Omicron

**DOI:** 10.1101/2022.09.25.509344

**Authors:** Xiangyang Chi, Lingyun Xia, Guanying Zhang, Ximin Chi, Bangdong Huang, Yuanyuan Zhang, Zhengshan Chen, Jin Han, Liushu Wu, Zeya Li, Hancong Sun, Ping Huang, Changming Yu, Wei Chen, Qiang Zhou

## Abstract

The pandemic of COVID-19 caused by SARS-CoV-2 continues to spread around the world. Mutant strains of SARS-CoV-2 are constantly emerging. At present, Omicron variants have become mainstream. In this work, we carried out a systematic and comprehensive analysis of the reported spike protein antibodies, counting the antibodies’ epitopes and genotypes. We further comprehensively analyzed the impact of Omicron mutations on antibody epitopes and classified these antibodies according to their binding patterns. We found that the epitopes of one class of antibodies were significantly less affected by Omicron mutations than other classes. Binding and virus neutralization experiments show that such antibodies can effectively inhibit the immune escape of Omicron. Cryo-EM results show that this class of antibodies utilizes a conserved mechanism to neutralize SARS-CoV-2. Our results greatly help us deeply understand the impact of Omicron mutations. At the same time, it also provides guidance and insights for developing Omicron antibodies and vaccines.

## Introduction

The pandemic of coronavirus disease 2019 (COVID-19) caused by severe acute respiratory syndrome coronavirus 2 (SARS-CoV-2) has lasted for nearly three years^1,2^. The spike (S) protein on the surface of the virus particle is the key protein for the virus to invade cells^3,4^. The S protein is a trimer containing multiple domains, of which the domain that directly binds to the receptor angiotensin-converting enzyme 2 (ACE2) is called the receptor binding domain (RBD)^3-5^. The RNA genome of SARS-CoV-2 is prone to mutations in the replication process, resulting in the constant emergence of mutant strains^6^. So far, several mutant strains have been identified as mutants worthy of attention by the World Health Organization, including Omicron and the previous Alpha^7^, Beta^8^, Gamma^9^ and Delta^10^ mutants. Among these mutant strains, Omicron contains the largest number of mutations and has stronger transmissibility than other mutant strains^11,12^. Omicron strains include several subtypes, such as BA.1-BA.5^13,14^. Mutations in the S protein confer stronger ACE2 affinity and immune escape ability^15-19^. Among them, there are 30-36 mutations located in the S protein, including 15-17 on the RBD. Some of these mutations can enhance the binding of the virus and the receptor, resulting in stronger viral infectivity^19,20^. Some other mutations can change the immunogenicity of the virus and give the virus the ability to escape^11,21^. This makes Omicron strain, especially BA.5, quickly replace the original prevalent strain and cause rapid and widespread transmission in the population^14^.

The neutralizing antibody is an important protective barrier against viral infection^22,23^. Antibodies against SARS-CoV-2 can be divided into RBD antibodies, N-terminal domain (NTD) antibodies and other antibodies according to their action sites^24-26^. These antibodies can also be identified as ordinary antibodies or nanobodies according to their types^25,26^. The complex structure of many antibodies with the viral S protein or RBD domain has been resolved^5,27-29^. The S proteins in these complexes are diverse, including wild-type and various mutant proteins. A comprehensive and systematic analysis of the epitopes and modes of action of these antibodies can help us deeply understand the working mechanism of antibodies.

In order to study the immune escape of Omicron in more detail, we comprehensively and systematically studied the interaction between the antibodies reported in PDB and current Omicron strains. Our results show that Omicron mutations affect the epitopes of most of the existing antibodies in PDB. Based on the binding mode of antibodies, we classified these antibodies and found that the epitopes of one class of antibodies were significantly less affected by Omicron mutations than other classes. Binding experiments and neutralization experiments showed that such antibodies could effectively inhibit the immune escape of Omicron. In addition, antibodies developed for Omicron BA.1 strain can effectively inhibit the other Omicron subtypes. Our work provides important insights into developing antibodies and a new generation of vaccines.

## Results

### Analysis of antibodies

We chose the antibodies of which complex structures with the S protein of SARS-CoV-2 have been resolved (Table 1). We found 518 complex structures of the antibody of SARS-CoV-2 with the S protein from the PDB database. Most of these complexes contain only one antibody (430, accounting for 83.01%), and the rest contain multiple antibodies as a cocktail combination. There are 82 complexes containing two antibodies (accounting for 15.83%), 5 complexes containing three antibodies (accounting for 0.97%) and 1 complex containing four antibodies (accounting for 0.19%) (Fig. 1A). In order to analyze the interaction between the antibody and the S protein in detail, we extracted subcomplexes from these complex structures. Each subcomplex contains an ordinary antibody or nanobody and its binding domain in S protein. A complex structure may contain multiple subcomplexes. A total of 613 subcomplex structures were obtained. Among them, 514 subcomplexes bound to the S protein of wild-type SARS-CoV-2, accounting for 83.85 % of the total, followed by Beta, Omicron, Delta, Kappa, Alpha and Gamma, with the number of antibodies being 44, 33, 8, 6, 3, 3 and 2 respectively (Fig. 1B). There is still a huge demand for developing antibodies against mutant strains such as Omicron.

**Fig. 1.**
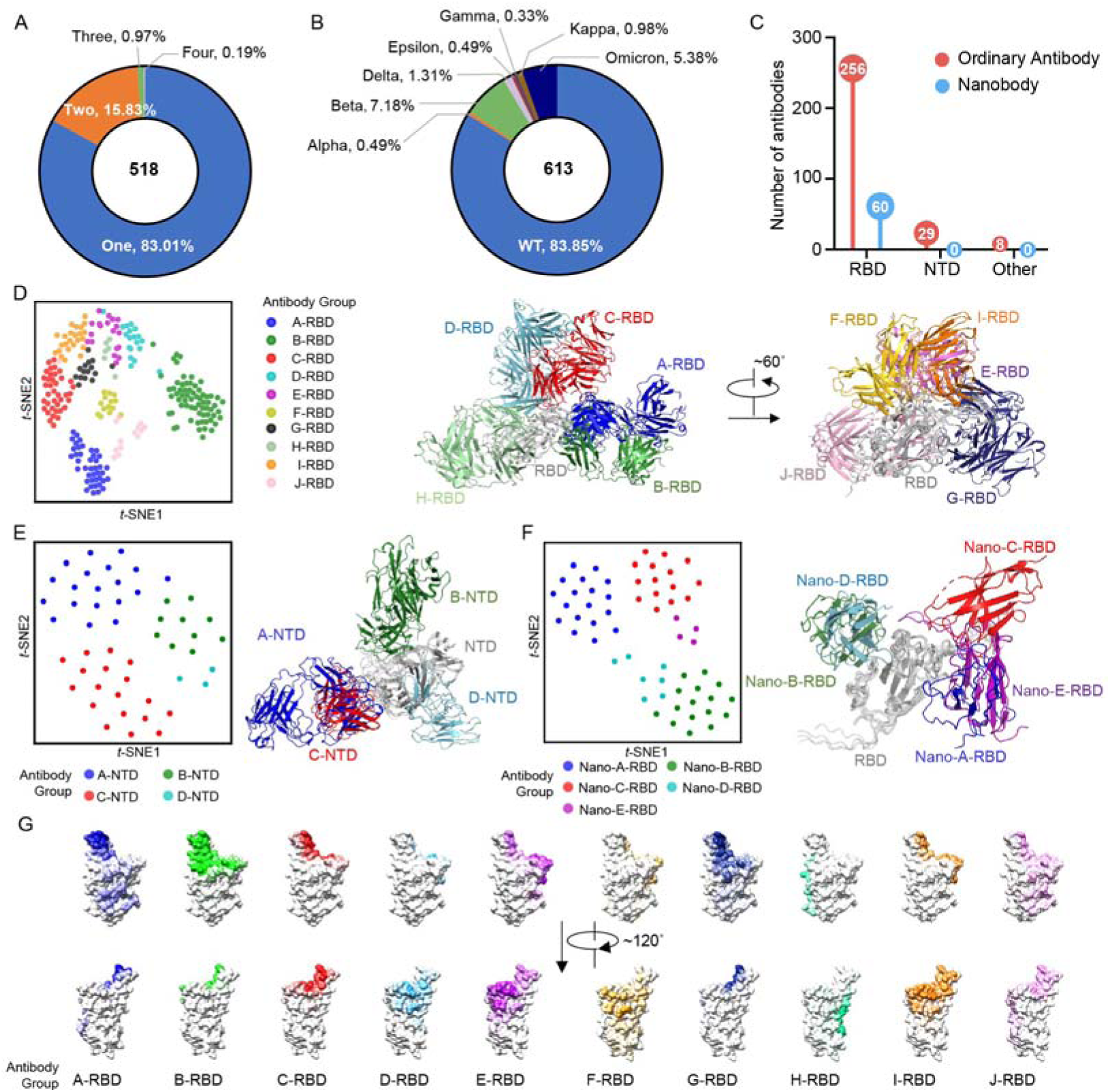
Antibody landscape of the structure available antibodies against SARS-Cov-2 S protein. (**A**) Distribution of the antibody number in the available structures. The antibody and corresponding binding domain are extracted as a subcomplex. Most structures contain only one antibody. (**B**) Strain distribution of the S protein in subcomplexes. Most of the S proteins are WT. (**C**) Statistics of ordinary antibodies and nanobodies and their corresponding targeting domains in the S protein. (**D**) Cluster distribution of the 10 classes of ordinary antibodies targeting RBD, named Ab-A-RBD to Ab-J-RBD, where Ab represents antibody and can be omitted as A-RBD to J-RBD without causing confusion, and structural scheme of the structural classes of some representative RBD antibodies. (**E)** are same as **D** for 4 classes of NTD antibodies (named Ab-A-NTD to Ab-D-NTD, where Ab represents antibody and can be omitted as A-NTD to D-NTD without causing confusion). (**F)** are same as **D** for the 5 classes of RBD nanobodies. For the t-SNE analysis of NTD antibodies, all of subcomplexes were used for analysis. For RBD antibodies, only the representative subcomplexes were used. (**G**) Epitope distributions of 10 classes of ordinary antibodies targeting RBD. The color depth represents the frequency of the residues as epitopes. The PDB ID for RBD of the S protein is 7QUS.

Considering that some antibodies contained in the subcomplex structures are duplicated, after removing these duplicated antibodies, we found a total of 293 ordinary antibodies and 60 nanobodies. Among them, there are 233 ordinary antibodies or nanobodies, each corresponding to only one subcomplex. Antibodies with multiple subcomplexes indicate that have attracted more research interest than other antibodies. Two antibodies, CR3022 and EY6A, have the largest number of subcomplexes at present (13 subcomplexes) (Fig. S1A). Among them, 5 single chain variable fragments (scFv) are included in the statistics when counting the number of epitope residues mutated in Omicron (NERMO) but not included in the genotypes. Among 293 ordinary antibodies, 256 bind the RBD domain, 29 bind the NTD domain, and 8 bind the other regions of the S protein. While 60 kinds of nanobody bind to RBD, there is a lack of nanobodies that bind to NTD (Fig. 1C).

We analyzed the genotype of the antibodies. From the perspective of genotype, the heavy chains of antibodies are mainly encoded by IGHV3, accounting for about half of the number of heavy chains. The numbers of antibodies coded by IGHV1 and IGHV4 are also relatively large, accounting for 29.97% and 11.50%, respectively. The light chains of antibodies are mainly encoded by IGKV1, IGKV3, IGLV2 and IGLV1, accounting for more than 75% (Fig. S1B). Due to the diversity of specific sources of nanobodies, we did not conduct genotype analysis and statistics of nanobodies.

We classified antibodies according to their spatial positions of binding RBD or NTD by comparing the structures of the subcomplexes (Fig. 1D-G and Fig. S1C-E). In order to retain epitope information to the maximum extent, we selected the subcomplex with the most complete structure of the S protein as the representative for analysis. The ordinary antibodies binding to RBD can be divided into 10 categories (Fig. 1D), named Ab-A-RBD to Ab-J-RBD, where Ab represents antibody and can be omitted as A-RBD to J-RBD without causing confusion. These antibodies covered almost all of the RBD surface, where B-RBD accounts for 25.39%, A-RBD accounts for 16.02%, and C-RBD class accounts for 14.84%, the three types of RBD antibodies with the largest proportion. We showed the epitopes of different antibodies structurally (Fig. 1G). The epitope patterns of A-RBD, B-RBD, and G-RBD are similar, while the epitope patterns of C-RBD, E-RBD, and I-RBD are relatively concentrated and distributed on the other side. All of these six classes of antibodies can interact with the antiparallel beta-sheet on RBD. The epitopes of the remaining four classes of antibodies are far away from the antiparallel beta-sheet, where D-RBD and H-RBD bind the loop region of R454-L492, F-RBD binds to the loop region of N437-W449, and J-RBD binds to the loop region of W495-P507. The structural comparison found that all 10 classes of RBD antibodies can bind to the up conformation of RBD. A-RBD, B-RBD, G-RBD, H-RBD, and J-RBD may cause steric hindrance to other protomers of S protein or NTD domain when RBD is in the down conformation (Fig. 1D, Fig. S1C).

Ordinary antibodies binding NTD can be divided into four categories, named Ab-A-NTD to Ab-D-NTD, where Ab represents antibody and can be omitted as A-NTD to D-NTD without causing confusion. Because the number of NTD antibodies is small, all of subcomplexes were used for analysis rather than using the representative subcomplex only (Fig. 1E). Based on the structure information, there were five loops in NTD, designated N1 (residues 14 to 26), N2 (residues 67 to 79), N3 (residues 141 to 156), N4 (residues 177 to 186), and N5 (residues 246 to 260). These antibodies bind to different regions of NTD (Fig. 1E, Fig. S2A), and the largest number of class, A-NTD, accounted for 41.38% of the total NTD antibodies. B-NTD class is the only class of antibodies binding the interface without an N-glycosylation site. Its epitope is close to the N2 region, accounting for 27.59%. C-NTD accounted for 24.14% of the total NTD antibodies. A-NTD and C-NTD classes bind to the N3 region of NTD, and D-NTD class mainly binds to the N4 loop (Fig. 1E, Fig. S1D).

The nanobodies against RBD can be divided into five categories (Fig. 1F), named Nano-A-RBD to Nano-E-RBD, where nano represents nanobody. Nano-A-RBD, Nano-C-RBD, and Nano-E-RBD classes are all bound to the receptor binding motif (RBM) region, accounting for 61.67% (Fig. S1E, Fig. S2B). For nanobodies, the RBM region is a hot area for antibody research and development. And the binding of these three kinds of nanobodies is not affected by RBD conformation. The other two types of Nano-B-RBD and Nano-D-RBD are bound in the core region of RBD, and when RBD is deeply closed, it will collide with the RBD domain of the adjacent protomer in space.

We further analyzed the genotypes corresponding to the heavy and the light chains of the antibodies of various structural types. Some genotypes can encode multiple structural types of heavy and light chains, such as IGHV3-30, which encodes 10 structural types of antibody heavy chains, and IGLV2-14, which encodes 9 structural types of antibody light chains, both of which cover RBD and NTD antibodies. While some genotypes encode only one structural type, for example, IGHV2-70 encodes only the heavy chain of J-RBD, and IGKV3D20 encodes only the light chain of A-RBD (Fig. S3A-B). From the perspective of structural types, some structural types have obvious genotype preferences, such as the heavy chain of B-RBD antibodies is mainly encoded by IGHV3-53 and IGHV3-66, and the light chain is mainly encoded by IGKV1-9, IGKV3-20 and IGKV1-33. Some classes have genotype preference only for heavy chains, such as the heavy chain of A-NTD is mainly encoded by IGHV1-24, and the light chain has no obvious genotype preference. Some heavy chain and light chain classes lack genotype preference, such as E-RBD, J-RBD and D-NTD, *etc*. From the perspective of genotype, some genotypes preferentially encode some structural types, such as IGHV3-53, which is mainly involved in the heavy chain coding of B-RBD and C-RBD antibodies, while IGKV3-20 is mainly involved in the light chain coding of B-RBD, A-RBD, and I-RBD (Fig. S3C).

### Effects of Omicron mutation

We analyzed the average number of epitope residues mutated in Omicron (ANERMO), including BA.1, BA.2, BA.3, BA.4, BA.5 and BA.2.12.1 subtypes, of all 353 ordinary antibodies or nanobodies. We counted the ANERMO of these antibodies. There are 44 antibodies with the ANERMO less than 0.5, accounting for 12.46% of the total. The ANERMO is greater than 2 for most antibodies (229, accounting for 64.87%), which indicates that most antibodies cannot inhibit immune evasion of Omicron (Fig. S4A). We counted and analyzed the strain type of the S protein in the 613 subcomplexes and found that antibodies developed against Omicron were least affected by Omicron mutation (the ANERMO was about 0.43), and epitopes of antibodies developed against Kappa are less affected by Omicron mutation than the other strains. However, the antibodies developed against Alpha, WT, Gamma and Beta strains have a relatively higher ANERMO (Fig. 2A).

**Fig. 2.**
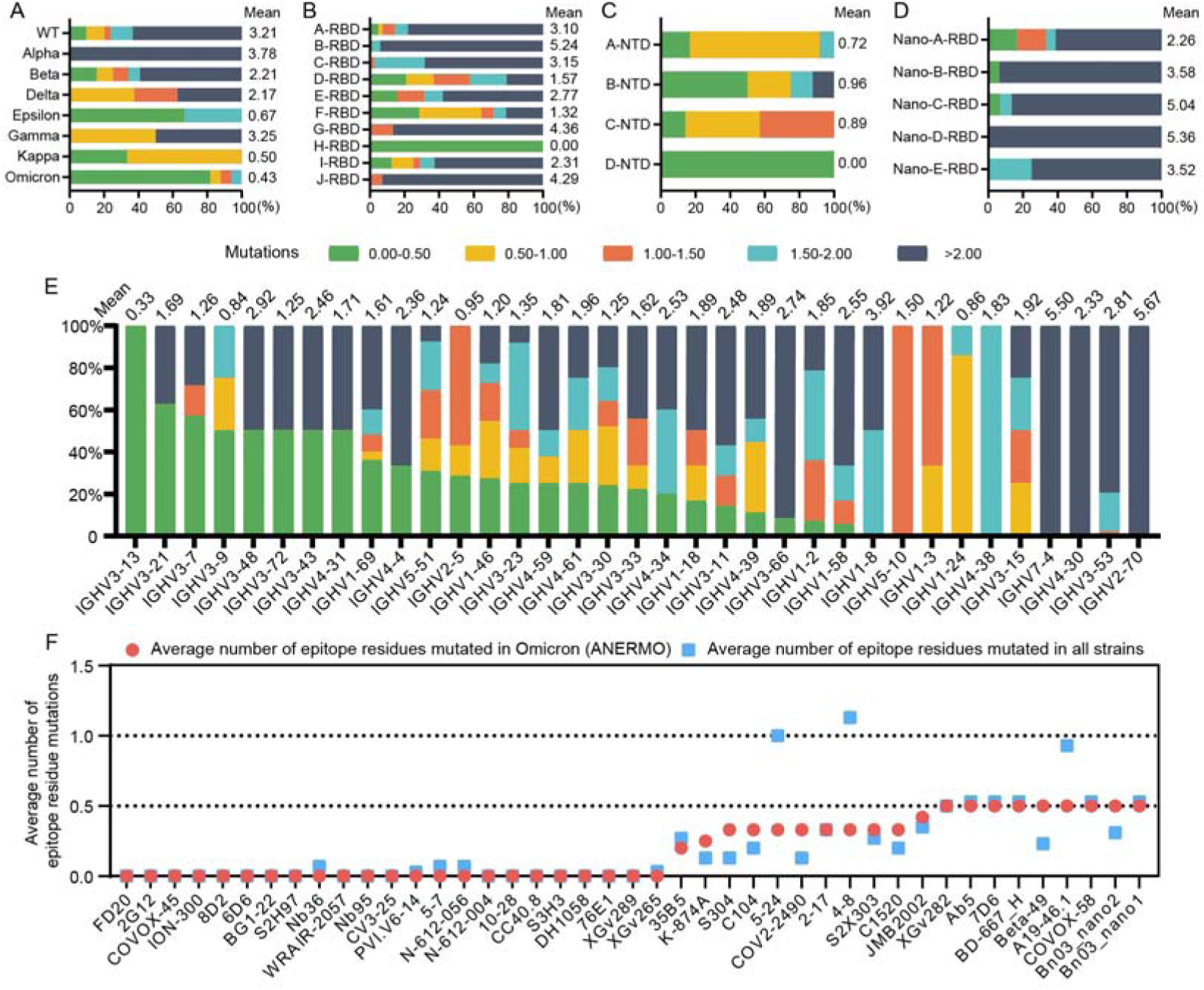
Epitope mutation statistics in Omicron. (**A**) Epitope mutations in Omicron of antibodies targeting different strains. The average number of epitope residues mutated in Omicron (ANERMO) of the antibodies targeting each strain are listed at the top of the columns. (**B**) The ANERMO from different RBD antibody structural classes. (**C**) The ANERMO from different NTD antibody structural classes. (**D**) The ANERMO from different RBD nanoantibody structural classes. (**E**) The ANERMO from different heavy chain genotypes. (**F**) Statistics of the potential broad-spectrum antibodies. Red points stand for ANERMO; blue squares stand for average number of epitope residues mutated in WT, Alpha, Lambda, Beta, Delta, Kappa, Gamma, and Omicron strains.

The extent affected by Omicron mutation varies for antibodies of different structural classes. The ANERMO of ordinary antibodies targeting RBD is 3.47, but it varies for different structural classes. The ANERMO of the H-RBD class is 0, significantly less than other types of antibodies (Fig. 2B, Fig. S4B-S4D). The ANERMO of B-RBD antibodies reached 5.24 (Fig. 2B).

NTD antibodies are less affected than RBD antibodies in general (Fig. 2B-C, Fig. S4B). The ANERMO of all NTD antibodies is 0.78. The D-NTD antibodies have no mutated epitopes in Omicron strains, which binds the N4 domain and may be a good target for developing the Omicron neutralizing antibodies (Fig. 2C, S4B, S4E). It is worth noting that the B-NTD class contains antibodies that bind to two infection-enhancing sites. These antibodies can help the RBD domain maintain an open conformation, which facilitates the receptor binding of SARS-CoV-2. The RBD nanobodies are generally more affected by Omicron mutation than ordinary antibodies (Fig. 2D), and the ANERMO of the RBD nanobodies is 3.94 (Fig. S4B, S4F).

From the perspective of antibody genotype, the heavy chains encoded by IGHV3-13, IGHV3-9 and IGHV1-24 are least affected by Omicron mutation with the ANERMO less than 1 (Fig. 2E). The heavy chains encoded by IGHV2-70 are greatly affected by Omicron mutation. The light chains are generally less affected by Omicron mutation than the heavy chains, while the light chains are less involved in binding to the S protein. In the term of the light chain genotypes, the antibodies encoded by IGLV8-61, IGKV2-24 and IGLV3-1 are least affected by Omicron mutation, while the epitopes of the antibodies encoded by IGKV1-9, IGLV5-37 and IGKV9-49 are mutated more in Omicron strains (Fig. S3D), which may not be able to effectively inhibit Omicron immune evasion.

### Evaluation of neutralization ability of potential broad-spectrum antibodies

We selected antibodies with few ANERMO (ANERMO <= 0.5) as “potential broad-spectrum antibodies” for further research. These potential broad-spectrum antibodies with few epitope residue mutations are likely to be able to maintain their binding ability and neutralize activity against Omicron. A total of 43 potential broad-spectrum ordinary antibodies or nanobodies were selected (38 ordinary antibodies and 5 nanobodies) (Fig. 2F). There were 23 kinds of antibodies whose epitope residues were not affected by Omicron mutations (ANERMO = 0), accounting for 6.5% of the total number of antibodies. Of the 43 potential broad-spectrum antibodies, 9 antibodies target the NTD, 28 antibodies are bound to the RBD, and the remaining 6 antibodies are bound to the S2 region. The total number of antibodies bound to the S2 region enrolled in this work is 7, which indicates that the antibodies developed against the S2 region are more likely to be used as broad-spectrum antibody drugs to avoid the immune escape problem caused by Omicron mutation.

The proportion of NTD antibodies among potential broad-spectrum antibodies (20.93%) is higher than that of NTD antibodies among all antibodies (8.22%) with the ANERMO of 0.22. We also selected antibodies with more affected epitope residues (ANERMO > 0.5), including 4-18 and CR3022.

We synthesized the gene of some representative potential broad-spectrum antibodies from different groups and some antibodies with high ANERMO as control and expressed these antibodies. We first determined the binding abilities of these antibodies to the S proteins of WT, Delta strain and Omicron subtypes. Among the 38 expressed antibodies, there were 12 antibodies whose EC_50_ values of binding to the S protein of WT, Delta strain or Omicron strain remained within the same range. 12 antibodies whose binding ability to one or two Omicron subtypes was at least one order of magnitude lower than that of WT, and 13 antibodies whose binding ability to three to five Omicron subtypes was at least one order of magnitude lower than that of WT. The binding ability of antibody 2G12 to Delta strain or Omicron subtypes was stronger than that of WT strain (Fig. 3A). In E-RBD, H-RBD, I-RBD, and S2 groups, there are half antibodies whose binding ability to S protein was not affected by Delta or Omicron mutation (Fig. 3A). Antibodies with less ANERMO and good broad-spectrum binding ability were mainly from the E-RBD, F-RBD, H-RBD and S2 groups (Fig. 3B).

**Fig. 3.**
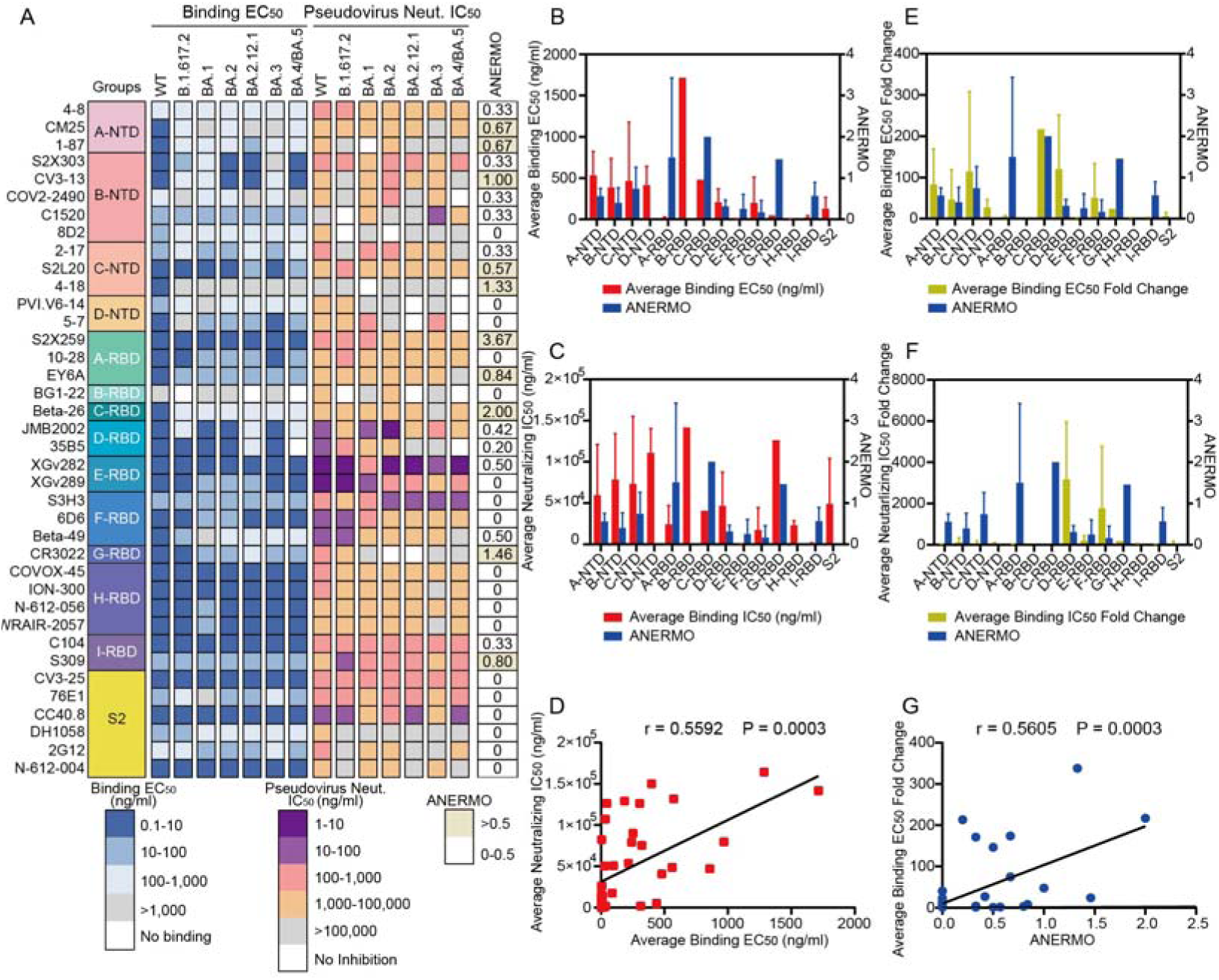
Binding and neutralization activities of the potential broad-spectrum antibodies. (**A**) Heatmap showing the binding EC_50_ against S fragments of SARS-CoV-2 variants and neutralization IC_50_ of pseudotyped SARS-CoV-2 variants. (**B**) The average binding EC_50_ against the WT, Delta, and the five indicated Omicron variants, and the average number of epitope residues mutated in Omicron (ANERMO) of different classes of antibodies. (**C**) The average neutralization IC_50_ against the WT, Delta, and the five indicated Omicron variants, and the ANERMO of different classes of antibodies. (**D**) The Pearson correlation coefficients analysis between the average binding EC_50_ and the average neutralization IC_50_ values of the expressed antibodies. (**E**) The average binding EC_50_ fold change between the five indicated Omicron variants and WT, and the ANERMO of different classes of antibodies. (**F**) The average neutralization IC_50_ fold change between the five indicated Omicron variants and WT, and the ANERMO of different classes of antibodies. (**G**) The Pearson correlation coefficients analysis between the average binding EC50 fold change and the ANERMO of the expressed antibodies. Samples were tested in triplicate. Data are shown as mean ± SD.

We measured the neutralization activity of these antibodies against WT, Delta, and Omicron pseudoviruses. The neutralization IC_50_ against WT, Delta and Omicron subtypes was less than 100 µg/ml for 17 antibodies (Fig. 3A), and the antibodies with high average neutralization activity and with less ANERMO were mainly from the E-RBD, F-RBD, H-RBD classes (Fig. 3C). There is a statistical correlation between the average IC_50_ of neutralizing activity of antibodies against Omicron subtypes and the average EC_50_ of these antibodies binding Omicron (*r* = 0.5592, *P* = 0.0003) (Fig. 3D), indicating that antibodies with a good binding ability to mutant strains often have good neutralizing activity against mutant strains.

We calculated the average fold change between the EC_50_ bound to Omicron and that to the WT (average EC_50_ fold change) and the average fold change between the IC_50_ of the antibodies to Omicron and that to WT (average IC_50_ fold change) for the 38 expressed antibodies. The results show that the E-RBD, H-RBD, and S2 groups have a low ANERMO (less than 0.5), low average EC_50_ change fold change, and low average IC_50_ fold change, indicating that the antibodies in these three groups were good broad-spectrum antibodies against a variety of Omicron strains (Fig. 3E and 3F). We calculated the relationship between the average EC_50_ change fold change and the ANERMO for the antibodies with ANERMO less than 3 and found a significant correlation between them (*r* = 0.5605, *P* = 0.0003) (Fig. 3G). Interestingly, the ANERMO of S2X259 was high (ANERMO = 3.67); however, it can maintain good binding and neutralizing activity against Omicron, which shows that our calculation method can effectively predict broad-spectrum antibodies with low ANERMO.

### Structural Analysis using cryo-EM

To further investigate the mechanism of these antibodies, we selected several antibodies with conserved epitopes, including A-RBD class antibody 10-28, E-RBD class antibodies XGv282 and XGv289, H-RBD class antibody COVOX-45, ION-300, S2H97, WRAIR-2057 and N-612-056, C-NTD class antibody S2L20, and S2 antibody CV3-25, and tried to solve the cryo-EM structures of these antibodies in complex with the S protein of Omicron BA.5. We have successfully solved the cryo-EM complexes of antibodies XGv282, XGv289, and S2L20 with the S protein in high-resolution (Figs. 4, S5-6, Table S1).

**Fig. 4.**
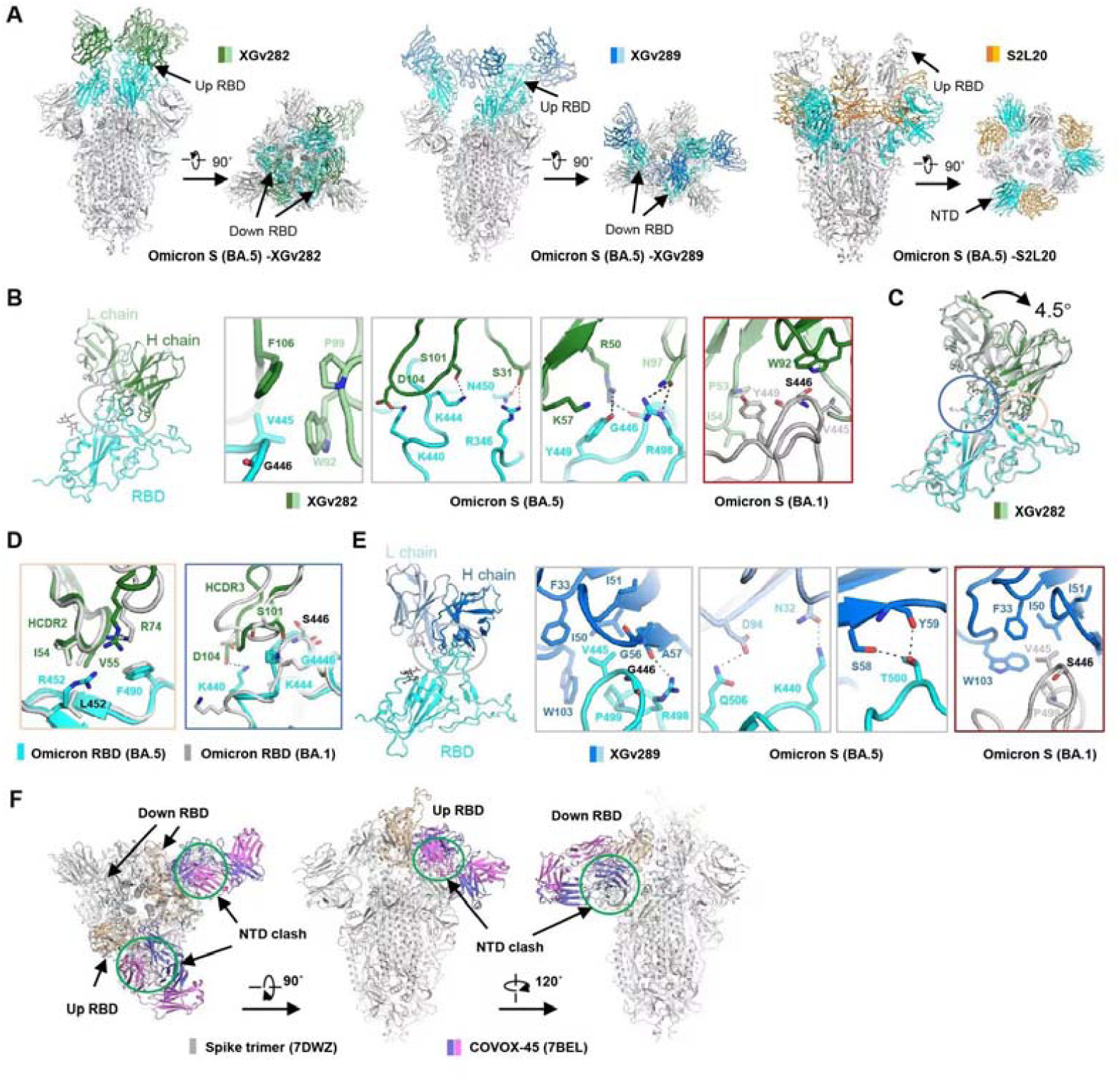
Structure analysis of Omicron BA.5 S protein in complex with antibodies. (**A**) The domain-colored cryo-EM structures of Omicron BA.5 S protein in complex with XGv282, XGv289 and S2L20 are shown in two perpendicular views. The domains (RBD or NTD) bound with antibodies are colored cyan, with the other part of the S protein shown in grey. The heavy and light chains are colored green and palegreen for XGv282, marine and blue for XGv289, and orange and yellow for S2L20, respectively. (**B**) The binding interface between Omicron BA.5 RBD and XGv282. Extensive hydrophobic and hydrophilic interactions on the interface are shown in three grey boxes. The red-framed right panel is the binding interface between Omicron BA.1 RBD and XGv282 (PDB ID: 7WLC). Black dashed lines indicate polar interactions. (**C**) Structural comparison of the Omicron BA.5 RBD-XGv282 complex and the Omicron BA.1 RBD-XGv282 complex (PDB ID: 7WLC). The RBD is superimposed, with the circled regions shown in detail in (**D**) The heavy and light chains of XGv282 are colored green and pale green, respectively. RBD is colored cyan and grey for Omicron BA.5 and BA.1, respectively. (**E)** is same as **B**, but for XGv289. The red-framed right panel is the binding interface between Omicron BA.1 RBD and XGv289 (PDB ID: 7WE9). (**F**) The RBD/COVOX-45 complex is superimposed on the RBD of the trimeric S protein (PDB ID: 7DWZ). COVOX-45 clashes with trimeric S protein in both the “up” and “down” conformation of RBD.

### E-RBD class antibodies XGv282 and XGv289

Both XGv282 and XGv289 have a strong neutralizing ability against each Omicron variant, the WT and Delta variants (Fig. 3A). The neutralization ability of XGv282 to BA.2 and BA.4/BA.5 is more potent than that to BA.1 and BA.3. And the neutralization ability of XGv289 to BA.1 was stronger than that to other strains. The XGv282 antibody binds to the BA.1 S protein through hydrogen bonds, cationic π bonds and numerous hydrophobic interactions^30^ (Fig. 4B). Compared with the XGv282/BA.1 S protein complex, the XGv282 antibody bound to the BA.5 S protein was rotated by 4.5° (Fig. 4C), which increased the interaction interface area between the antibody and the S protein (the interface area of XGv282 with BA.5 or BA.1 S protein is 750 Å^2^ or 550 Å^2^, respectively) (Fig. S7A). The L452R mutation exists in the BA.5 S protein, considered to evade HLA-A24-restricted cellular immunity^31^, which disrupted the hydrophobic interaction between XGv282 and the BA.5 S protein (Fig. S7B). Additionally, the residue F490 shifted 3.0 Å further away from R74 with respect to the native structure (XGv282/BA.1 S protein), which affects the binding ability between XGv282 and S protein (Fig. 4D). The mutation of the critical residue S446G of the BA.5 S protein (Ser in BA.1 and BA.3) resulted in a conformational change in the loop region, reshaped the interaction network with XGv282 and formed seven pairs of hydrogen bonds (K444-S31, K444-S101, G446-R50, Y449-R50, Y449-K57, N450-S31 and K498-N97) and a pair of cationic π bond (Y449-K57) (Fig. 4B). Moreover, the everted K444 interacts with S101 on HCDR3 of XGv282, K440 forms a salt bridge with D104 on HCDR3, and the turning of the F106 side chain extends the hydrophobic interaction interface, all of which further stabilize the interaction between XGv282 and BA.5 S protein (Fig. 4B).

The epitopes of XGv289 on the BA.5 S protein are located in two loop regions (N439-G447 and F497-Q506) subject to high mutation (Fig. S8A). Similar to XGv282/BA.5 S protein, the loop containing S446G and L452R showed slight conformational changes, which resulted in the overall rotation of XGv289 by 9.1° (Fig. S8B). It increased its overall interface (the interface area of XGv289 with BA.1 or BA.5 S protein is 550 Å^2^ or 650 Å^2^, respectively) mainly through hydrogen bonds (Y501-S99, T500-S58, T500-Y59 and V503-G95) and hydrophobic interactions (Fig. S8A). The binding experiment also shows that the affinity of XGv289 to the BA.5 S protein is greater than that to the BA.1 S protein (Fig. 4E).

### H-RBD antibody utilizes a conserved mechanism to neutralize SARS-CoV-2

H-RBD is a class of antibodies that binds at the site of the RBD flank, with a high affinity for S proteins of the SARS-CoV-2 variants, such as WT, Delta, and Omicron variants (Fig. 3A). In our present work, six antibodies were classified into the H-RBD class, including FD20, COVOX-45, ION-300, S2H97, WRAIR-2057 and N-612-056. Such antibody binding sites are hidden away from the RBM, but FD20 can still inhibit the interaction of the RBD with the receptor ACE2^32^. In addition, the binding sites of FD20, COVOX-45, ION-300, WRAIR-2057 and N-612-056 are highly conserved among the mainstream strains that have appeared so far.

To further characterize the commonalities of these antibodies, we investigated five H-RBD antibodies using cryo-EM. We showed that these antibodies could induce the dissociation of the trimeric S protein after incubation with the S protein (Fig. S9), which is consistent with the previous report about the antibodies FD20^32^ or N-612-056^33^. In addition, the neutralization ability of N-612-056 was related to incubation time, which may be due to the poor accessibility of its targeted epitopes and its dependence on the conformational change of the S protein^33^. We also make a superposition between the H-RBD antibody/RBDs and the antibody/S protein trimer with different conformational structures from the PDB database (PDB numbers 6VSB and 7BEL, respectively), indicating the epitopes of H-RBD antibodies are hidden in the trimeric S protein, regardless of whether the RBD domain of the S protein is up or down. When the RBD is in the “down” state, the binding site of H-RBD antibodies is close to the NTD of the adjacent monomer, causing a significant steric hindrance between the antibodies (Fig. 4F). Even when the RBD is in the “up” state, the clash between the antibodies and the NTD, or other parts of the S protein, can still be observed. Thus, we suggested that the antibodies of the H-RBD class are more likely to inhibit virus infection by destabilizing the S protein and inducing S protein dissociation.

### C-NTD class antibody S2L20

Antibody S2L20, a member of the C-NTD class of antibodies, has a strong binding affinity for the S protein and broad neutralization efficiency against the WT, Delta and Omicron variants (Fig. 3A). In this work, we solved the cryo-EM structure of the S2L20/BA.5 S protein complex at 3.1 Å (Fig. S10A). Each subunit binds to one S2L20 antibody with a highly conserved epitope near the glycan flank at position N234 of NTD, close to the RBD side of the same monomer. And the interface was stabilized by three pairs of hydrogen bonds and two pairs of salt bridges (Fig. S10B), which are consistent with the previously reported structure of S2L20-binding Delta variant S protein^34-36^.

## Discussion

Extensive efforts have been made to classify antibodies targeting the S protein using various methods. The classification of ordinary antibodies or nanobodies described previously was based on the mutant profile of epitopes^37^ or whether they can block the binding with the ACE2 and recognize different states (up or down or both) of the RBD^38,39^ or biochemistry evidence of pairwise competition between antibodies^40,41^ or binding to different epitope bins^42-44^. The accumulation of 3D structures of the S protein in complex with antibody/nanobody allowed us to characterize a panel of over 300 antibodies/nanobodies based on a new, systematic, and unbiased method that classified antibodies or nanobodies utilizing the structures. There are several advantages of this method. Firstly, this method is explicit and accurate. Secondly, it requires no library screening for the mutant profile or competitive binding assay. Finally, the antibody distribution relative to the S protein is tightly linked to the epitope bins, so the classification of antibody based on structure information is also coupled with different types of epitope bins. This method allows for a more detailed classification of antibodies or nanobodies, which provides new insights into the classification of antibodies.

In this work, we obtained several antibodies with relatively conserved epitopes, mainly from E-RBD, H-RBD, and C-NTD classes. Among them, the E-RBD antibodies XGv289 and XGv282 have the most potent ability to neutralize the current mainstream Omicron BA.5 strain, as these two antibodies were initially developed for Omicron BA.1 strain. In addition, mutations of S446G and L452R in BA.5 remodeled the interactions between XGv282 and RBD, increasing the interaction surface and affinity. The H-RBD antibodies have a high affinity for S protein of various strains, including WT, Delta and Omicron variants, epitopes of which are highly conserved. Previous studies have revealed the epitopes of H-RBD at a cryptic site in the RBD. The binding of the H-RBD class antibodies will induce the trimeric S protein to dissociate due to the steric hindrance. This may explain why there are no complex structures of this class of antibodies with trimeric S protein reported.

We note some potential broad-spectrum antibodies with fewer ANERMO, such as 4-8, 5-24, and A19-46.1. However, when considering more strains, the number of epitope residue mutations of these antibodies is relatively large. Among them, 4-8 and 5-24 have a large number of epitope residue mutations in the C.37 strain, while A19-46.1 has epitope mutations in multiple non-Omicron strains.

S2X259 maintains good binding and neutralizing activity against Omicron strains, although the ANERMO is high (ANERMO > 3). Thus, mutation of epitope residues may increase or decrease affinity. This work does not predict the specific effects of epitope mutations but only analyzes and counts the number of epitope mutations. Among the antibodies expressed and verified in this work, the antibody with a number of epitope mutations less than 3 has a significant correlation between its activity change fold and the number of epitope mutations, indicating that our method is still reliable. To more accurately predict the effect of antibody epitope mutations and screen for broad-spectrum antibodies, it can calculate the changes of parameters such as surface binding Gibbs free energy based on analyzing the number of epitope mutations. In the follow-up work, a reliable method for predicting the effect of mutations on activity can be realized so that the broad-spectrum activity of antibodies with a high number of mutations can be predicted more accurately.

## Acknowledgments

We thank the cryo-EM facility and the High-Performance Computing Center of Westlake University for providing technical support. This research was supported by grants from the National Natural Science Foundation of China (projects 32022037), Hangzhou agricultural and social development scientific research project (202204B14), the Young Elite Scientists Sponsorship Program by CAST, the Leading Innovative and Entrepreneur Team Introduction Program of Hangzhou, the Special Research Program of Novel Coronavirus Pneumonia of Westlake University and Tencent Foundation.

## Author contributions

Q.Z., W.C., and C.Y. conceived and supervised the project. XY.C., L.X., G.Z., Y.Z., J.H., L.W., Z.L., H.S., and P.H. did the experiments. XM.C., B.H., and Z.C. analyzed the PDB structures. XY.C., L.X., G.Z., XM.C., B.H., Y.Z. and Z.C. performed the visualization. All authors contributed to the data analysis. XY.C., L.X., G.Z., XM.C., B.H., Y.Z., Z.C. and Q.Z. wrote the draft of the manuscript. Q.Z., W.C., and C.Y. reviewed and edited the manuscript. All authors have read and approved the article.

## Data availability

Atomic coordinates and cryo-EM density maps of the S protein of Omicron BA.5 SARS-CoV-2 in complex with antibodies (PDB ID: 8GTO, 8GTP and 8DTQ; EMDB ID: EMD-34259, EMD-34260, EMD-34261, EMD-34262, EMD-34263 and EMD-34264) have been deposited to the Protein Data Bank (http://www.rcsb.org) and the Electron Microscopy Data Bank (https://www.ebi.ac.uk/pdbe/emdb/),, respectively. Materials and data will be shared upon request.

## Supplementary Materials

Materials and Methods

Figs. S1 to S10

Table S1

