## Supplemental Materials for "Comprehensive structural analysis reveals broad-spectrum neutralizing antibodies against Omicron"

**This file includes:**

Materials and Methods

Figs. S1 to S10

Table S1

Supplemental References

**Materials and methods**

**Statistics and analysis of antibodies**

Structures of SARS-CoV-2 S protein in complex with antibody or nanobody were obtained from the Protein Data Bank (<https://www.rcsb.org/>). We labeled chains corresponding to the S protein and the associated heavy chain and light chain of antibody or nanobody manually in these structures. Subcomplexes were extracted from structures as separated PDB files utilizing UCSF Chimera. There may be more than one subcomplex corresponding to one antibody since multiple kinds of antibodies can exist in one structure.

Subsequently, the name and amino acid sequence of heavy chain and light chain of antibody or nanobody were obtained from EMBL-EBI (<https://www.ebi.ac.uk/pdbe/api/pdb/entry/status/>). Genotype analysis was performed using NCBI IgBlast (v1.17.1) with the IMGT reference (<https://www.ncbi.nlm.nih.gov/igblast/>) with default parameters except that the number of germline gene was set to 1. The strain type of the S protein of each structure was determined using sequence alignment.

Antigen epitopes were defined as residues that contact antibody within a distance of 4 Å in the spike protein. Subsequently, the binding domain of the antibody or nanobody was figured out using epitopes. By comparing the mutant profile of Omicron strains (BA.1, BA.2, BA.2.12.1, BA.3, BA.4/5) with the epitopes recognized by the antibody, the number of epitopes affected by Omicron mutations was computed. Noted that one antibody may form complexes with multiple strains of S protein resulting in multiple versions of epitopes under which condition the mean value was used.

To classify antibodies in three dimensions, the S proteins in the extracted subcomplexes were superimposed together firstly. And then the rotation angles among antibodies were calculated (for antibodies, only heavy chains were considered), resulting in an NxN matrix (N represents the number of antibodies), based on which the structural classes were computed with Python. Data were post-processed and visualized using Python.

**Generation and purification of monoclonal antibodies**

For ordinary antibodies, the sequences of their variable region were codon-optimized and synthesized in Azenta and then inserted into pcDNA3.4 vector plasmid containing human IgG1 heavy chain constant region sequence or light chain constant region sequence (κ or λ chain). To express these antibodies, 15 μg heavy chain plasmid and 15 μg light chain plasmid of the same antibody were cotransfected into 30 mL Expi293F cells using ExpiFectamine™ 293 Transfection Kit (Thermo Fisher) according to the manufacturer’s instructions. The Expi293F cells were then cultured for 120 hours in a shaker under 37°C and 5% CO_2_ condition. The expressed productions were centrifuged at 4000×g in 4°C for 20 minutes and the supernatants were filtered through a 0.22 μm filter. The filtered supernatant was then subjected to ProteinG column (Cytiva) which was equilibrated with phosphate buffered saline (PBS), pH 7.4. After washing with 5 column-volume (CV) PBS, the antibody was eluted with 5 CV 0.1 M Glycine, pH 2.7 and immediately neutralized to pH 7.4 using 1 M Tris-HCl, pH 9.0. The eluted antibody was then buffer exchanged into PBS and concentrated, pH 7.4 using 30 kDa Ultrafiltration centrifugal tube. The concentration of purified antibodies was measured using Nanodrop Plus (Thermo Fisher). The antibodies were then aliquoted and stored in -80°C until use.

**Enzyme linked immunosorbent assay (ELISA)**

The binding activities of purified antibodies to S protein of SARS-CoV-2 variants were tested using ELISA. Recombinant proteins of the extracellular domain of the S protein (S-ECD) of SARS-CoV-2 WT (Sino Biological, 40589-V08H4), B.1.617.2 (Sino Biological, 40589-V08B16), BA.1 (Acro Biosystems, SPN-C52Hz), BA.2 (Acro Biosystems, SPN-C5223), BA.2.12.1 (Acro Biosystems, SPN-C522d), BA.3 (Acro Biosystems, SPN-C5225), BA.4 (Acro Biosystems, SPN-C5229) were coated onto 96-well plates at the concentration of 2 μg/mL overnight at 4°C. The plates were washed three times with PBS plus 0.2% Tween (PBST) followed by incubation at 37°C for 1 hour with PBST containing 2% bovine serum albumin (BSA). Following washing with PBST, four-fold serial-diluted antibodies diluted in PBST containing 0.2% BSA starting at 1 μg/mL were added to the wells in triplicates and allowed to incubate at 37°C for 1 hour. After washing with PBST, horseradish peroxidase (HRP) conjugated anti-human IgG antibody (Abcam) diluted in PBST containing 0.2% BSA at the dilution of 1:10000 was added to each well and incubated at 37°C for 1 hour. After washing with PBST, TMB substrate solution (Solarbio) was added to the plates and incubated for 6 minutes at room temperature. ELISA stop solution (Solarbio) was added to stop the reaction. The absorbent at 450 nm was measured using microplate reader (TECAN) with the absorbent at 630 nm as reference. The data were processed using GraphPad Prism v8.3 and the EC_50_ values were calculated using a four-parameter nonlinear regression model.

**Neutralizing assay with pseudotyped SARS-CoV-2 variants**

The pseudotyped SARS-CoV-2 WT, B.1.617.2, BA.1 and BA.2 variants were packaged as previously described. Briefly, HIV backbone plasmid pNL4-3.Luc.R-E- was cotransfected into HEK293T cells with pCAGGS vector carrying full-length S protein gene sequence belonging to SARS-CoV-2 WT (QHD43416.1), B.1.617.2 (EPI_ISL_2029113), BA.1 (EPI_ISL_6640917) or BA.2 (YP_009724390.1) at different ratio, respectively. Supernatants containing pseudotyped virus were harvested at 48 hours, 60 hours, and 72 hours post-transfection, filtered through a 0.45-μm filter, aliquoted and stored at −80 °C. The pseudotyped viruses of SARS-CoV-2 BA.2.12.1 (DD1777), BA.3 (DD1774), BA.4 (DD1776) variants were purchased from Vazyme.

To determine the neutralizing efficacy of antibodies, 50 μL monoclonal antibody diluted in DEME medium containing 10% fetal bovine serum (FBS) (starting concentration at 100 μg/mL) was incubated with 50 μL pseudotyped virus in 96-well cell culture plates at 37 °C for 1 hour. ACE2-293T cells at the concentration of 2.5×10^5^ cells/mL in 100 μL DMEM medium supplemented with 10% FBS were then added into each well. The plates were incubated at 37°C and 5% CO_2_ condition for 48 hours. After incubation, 100 μL cell culture medium was discarded and 100 μL Brite-Lite Luciferase reagent (Vazyme) was added to each well and incubated for 2 minutes avoid from light. Each well was then mixed 10 times by pipetting, and 150 μL mixture was transferred to a white plate to measure the luciferase activity using a microplate reader (TECAN Spark). The inhibition percent was determined by comparing the relative luminescence units to the cell control (cells without pseudotyped virus or antibody) and virus control (cells with pseudotyped virus but without antibody). The data were processed using GraphPad Prism v8.3 and the IC_50_ values were calculated using a three-parameter nonlinear regression model.

**Protein expression and purification**

The S-ECD (1-1208 a.a) of Omicron BA.5 was cloned into the pCAG vector (Invitrogen) with substitution of six prolines at residues 817, 892, 899, 942, 986 and 987 ^1^, a “GSAS” substitution at residues 682 to 685 and a C-terminal T4 fibritin trimerization motif followed by one Flag tag. The recombinant S-ECD protein was overexpressed using HEK 293F mammalian cells (Invitrogen) at 37℃ under 5% CO_2_ in a Multitron-Pro shaker (Infors, 130 rpm). When the cell density reached 2.0 ×10^6^ cells/mL, the plasmid was transiently transfected into the cells. To transfect one liter of cell culture, about 1.5 mg of the plasmid was premixed with 3 mg of polyethylenimines (PEIs) (Polysciences) in 50 mL of fresh medium for 15 mins before adding to cell culture. Cells were removed by centrifugation at 4000×g for 15 mins after sixty hours transfection. The secreted S-ECD protein was purified using anti-FLAG M2 affinity resin (Sigma Aldrich). After loading two times, the anti-FLAG M2 resin was washed with the wash buffer containing 25 mM Tris (pH 8.0), 150 mM NaCl. The protein was eluted with the wash buffer plus 0.2 mg/mL Flag peptide. The eluent was then concentrated and subjected to size-exclusion chromatography (Superose 6 Increase 10/300 GL, GE Healthcare) in a buffer containing 25 mM Tris (pH 8.0), 150 mM NaCl. The peak fractions were collected and concentrated to incubate with antibody. The purified S-ECD was mixed with the antibody at a molar ratio of about 1:5 for one hour. Then the mixture was subjected to size-exclusion chromatography (Superose 6 Increase 10/300 GL, GE Healthcare) in buffer containing 25 mM Tris (pH 8.0), 150 mM NaCl. The peak fractions were collected for cryo-EM analysis.

**Cryo-EM sample preparation**

The peak fractions of complex were concentrated to about 3.5 mg/mL and applied to the grids. Aliquots (3.5 μL) of the protein complex were placed on glow-discharged holey carbon grids (Quantifoil Au R1.2/1.3). The grids were blotted for 3.5 s and flash-frozen in liquid ethane cooled by liquid nitrogen with Vitrobot (Mark IV, Thermo Scientific). The prepared grids were transferred to a Titan Krios operating at 300 kV equipped with Gatan K3 detector and GIF Quantum energy filter. Movie stacks were automatically collected using AutoEMation ^2^, with a slit width of 20 eV on the energy filter and a defocus range from -1.2 µm to -2.2 µm in super-resolution mode at a nominal magnification of 81,000×. Each stack was exposed for 2.56 s with an exposure time of 0.08 s per frame, resulting in a total of 32 frames per stack. The total dose rate was approximately 50 e-/Å2 for each stack.

**Data processing**

The movie stacks were motion corrected with MotionCor2 ^3^ and binned 2-fold, resulting in a pixel size of 1.077 Å/pixel. Meanwhile, dose weighting was performed ^4^. The defocus values were estimated with Gctf ^5^. Particles of Omicron BA.5 S in complex with XGv289 were automatically picked using Relion 3.0.6 ^6-9^ from manually selected micrographs. After 2D classification with Relion, good particles were selected and subject to 2D classification and multiple cycle of heterogeneous refinement without symmetry using cryoSPARC ^10^.The good particles were selected and subjected to non-uniform refinement, local CTF refinement and local refinement with C1 symmetry, resulting in the 3D reconstruction for the whole structures, which was further subject to local refinement with adapted mask on the interface between RBD of Omicron BA.5 S and XGv289 to improve the map quality on RBD-XGv289 subcomplex.

For Omicron BA.5 S (spike protein) in complex with XGv282 and S2L20, the CTF of stacks were estimated by patch CTF estimation (multi), and particles were automatically picked by template picker and extracted with cryoSPARC. The subsequent processing methods are consistent with the above description of Omicron BA.5 S in complex with XGv289.

The resolution was estimated with the gold-standard Fourier shell correlation 0.143 criterion ^11^ with high-resolution noise substitution ^12^. Refer to Fig. S5-S6 and Table S1 for details of data collection and processing.

**Model building and structure refinement**

For model building of Omicron BA.5 S in complex with XGv282, XGv289 and S2L20, the atomic model of the S-ECD (PDB ID: 7DWZ) were used as templates, which were molecular dynamics flexible fitted ^13^ into the whole cryo-EM map and manually adjusted with Coot ^14^ to obtain the atomic model of Omicron BA.5 S protein. The reported models of XGv282 (PDB ID: 7WLC), XGv289 (PDB ID: 7WE9) and S2L20 (PDB ID: 7SO9) were manually refined based on the focused-refined cryo-EM map. Each residue was manually checked with the chemical properties taken into consideration during model building. Several segments, whose corresponding densities were invisible, were not modeled. Structural refinement was performed in Phenix ^15^ with secondary structure and geometry restraints to prevent overfitting. To monitor the potential overfitting, the model was refined against one of the two independent half maps from the gold-standard 3D refinement approach. Then, the refined model was tested against the other map. Statistics associated with data collection, 3D reconstruction and model building were summarized in Table S1.

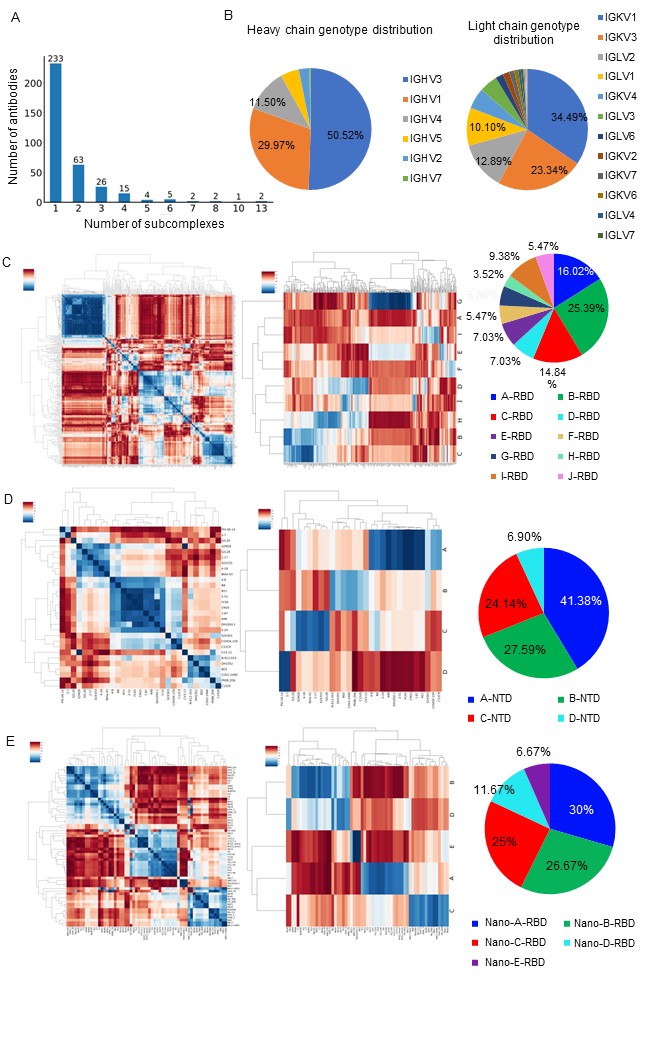
 **Fig. S1**

Statistics of the antibodies involved in this study.

**(A**) Statistics of the number of the subcomplexes corresponding to antibodies. (**B**) Genotype distribution of the heavy chains and the light chains. (**C**) Clustering results of RBD antibodies. On the left is cross-correlation analysis result. In the middle is class distribution. The ratio of different structural classes is depictured in the right. 10 classes of ordinary antibodies targeting RBD are named Ab-A-RBD to Ab-J-RBD, where Ab represents antibody and can be omitted as A-RBD to J-RBD without causing confusion. **(D)** and (**E)** are same to **C**, but for NTD antibodies and RBD nanobodies, respectively.

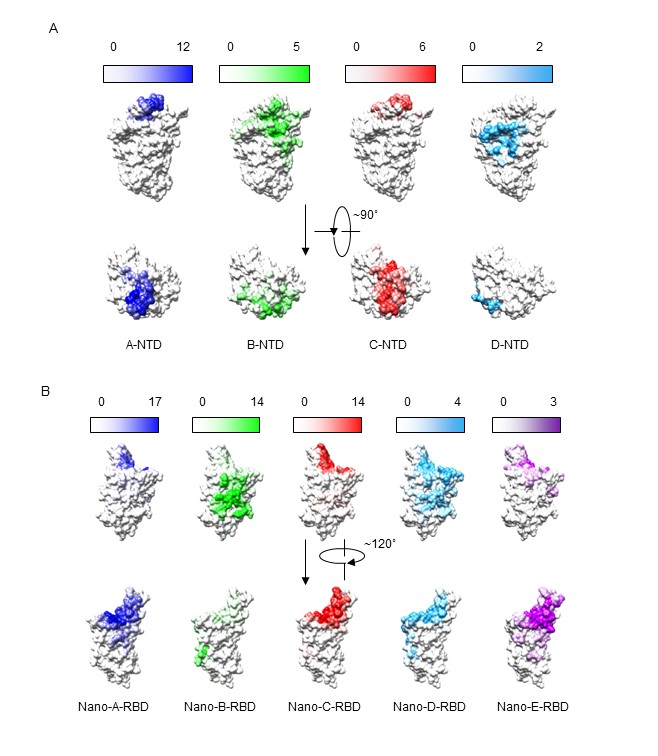

**Fig. S2**

Epitope distribution of different structural classes of antibodies.

**(A**) Epitope distribution of 4 classes of NTD antibodies. The color depth represents the frequency of the residues as epitopes. (**B)** is same as **A**, but for 5 classes of RBD nanobodies. The PDB ID for RBD or NTD of the S protein is 7QUS.

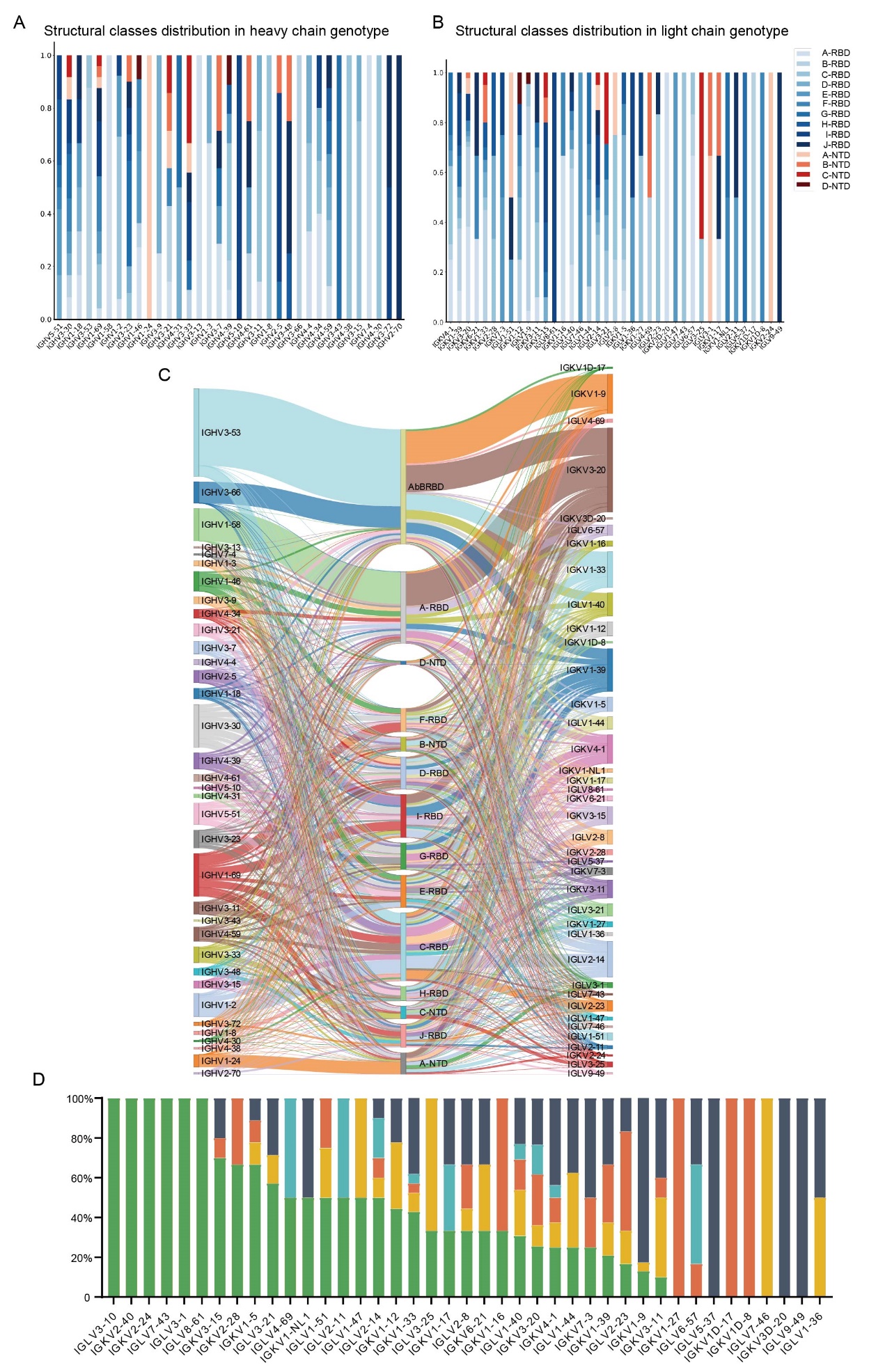

**Fig. S3**

Structural classes distribution verses chain genotypes.

(**A**) Structural classes distribution in heavy chain genotypes. (**B**) Structural classes distribution in light chain genotypes. (**C**) Corresponding relation of heavy chain genotypes, light chain genotypes and structural classes. (**D**) The average number of epitope residues mutated in Omicron (ANERMO) from different light chain genotypes.

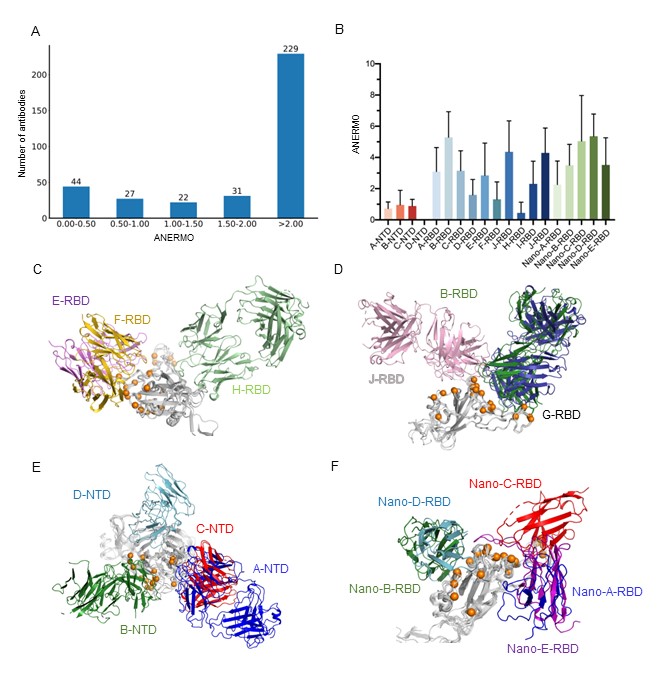

**Fig. S4**

Average epitope mutation numbers in Omicron.

(**A**) Distribution of average epitope mutation numbers in Omicron of antibodies. The average epitope mutation numbers are grouped in five (0.00-0.50, 0.50-1.00, 1.00-1.50, 1.50-2.00, and more than 2.00). The number of corresponding antibodies is shown on the top. (**B**) Epitope mutation number in Omicrons verses different structural classes with NTD antibodies in red, RBD antibodies in blue, and RBD nanobodies in green. (**C**) Structures of RBD antibody classes of E-RBD, F-RBD, and H-RBD. (**D**) Structures of RBD antibody classes of B-RBD, J-RBD, and G-RBD. (**E**) Structures of NTD antibody classes. The Cα of mutated residues in RBD are high-lighted in orange spheres. (**F**) Structures of RBD nanobody classes.

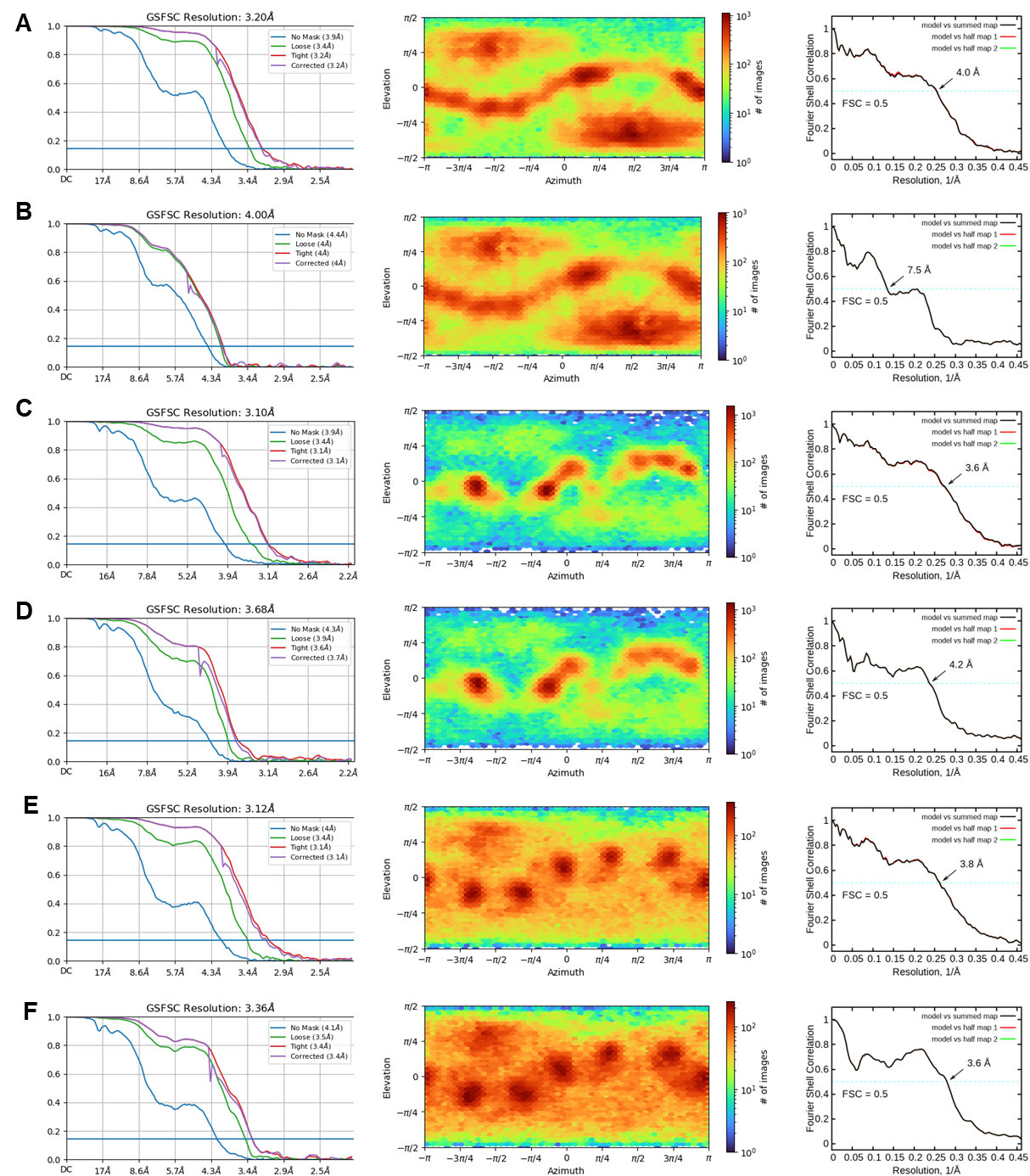

**Fig. S5**

Cryo-EM analysis of Omicron BA.5 S protein in complex with antibodies.

The left panel is the FSC curves. The middle panel is the Euler angle distribution. The right panel is FSC curve of the refined model versus the overall structure that it is refined against (black); of the model refined against the first half map versus the same map (red); and of the model refined against the first half map versus the second half map (green). The small difference between the red and green curves indicates that the refinement of the atomic coordinates is not enough overfitting. **(A-F)** are for Omicron BA.5 S protein in complex with XGv282, XGv289, S2L20, RBD of Omicron BA.5 in complex with XGv282, XGv289, NTD of Omicron BA.5 in complex with S2L20, respectively.

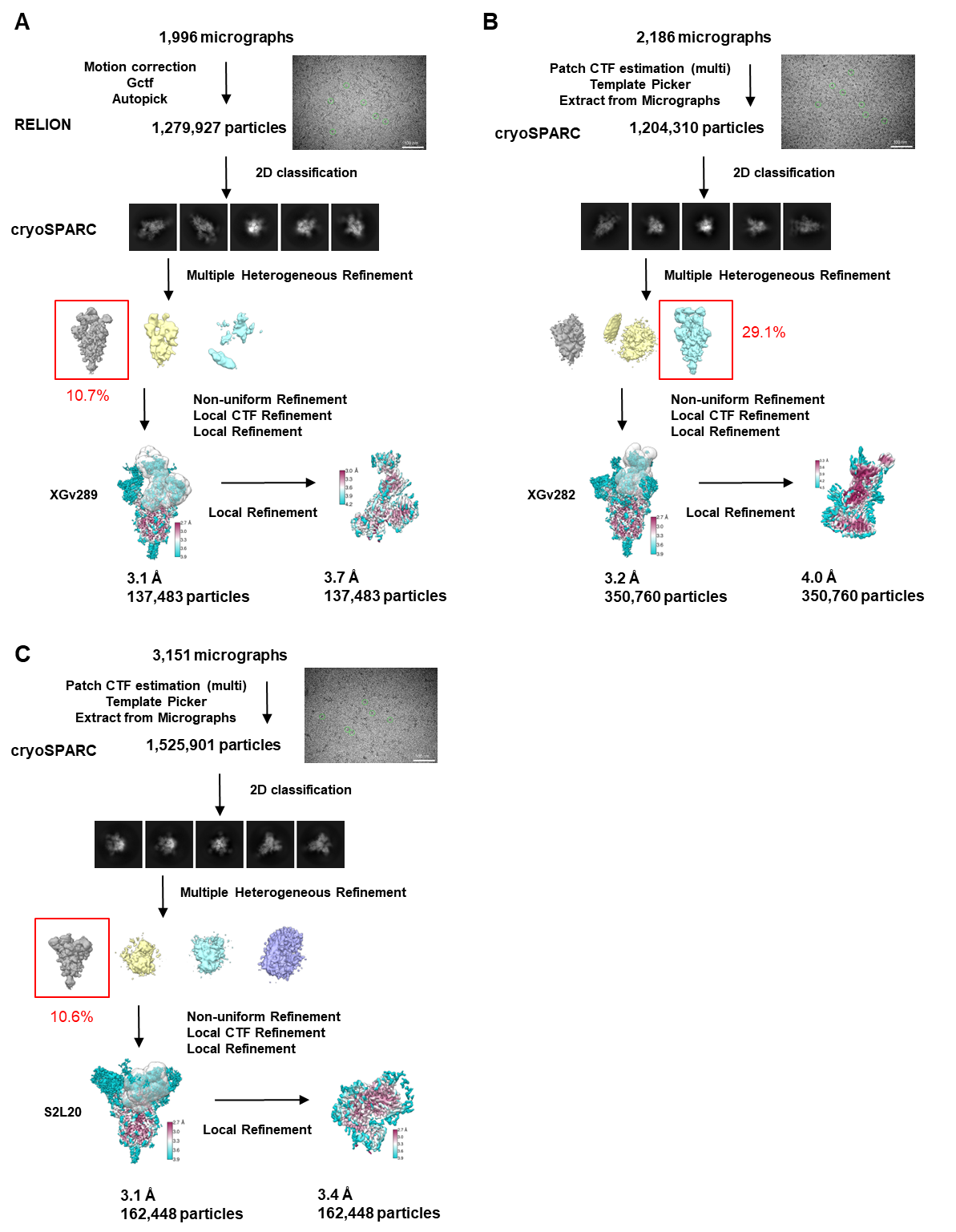

**Fig. S6**

Flowchart for cryo-EM data processing.

Please refer to the ‘Data Processing’ section in Methods for details.

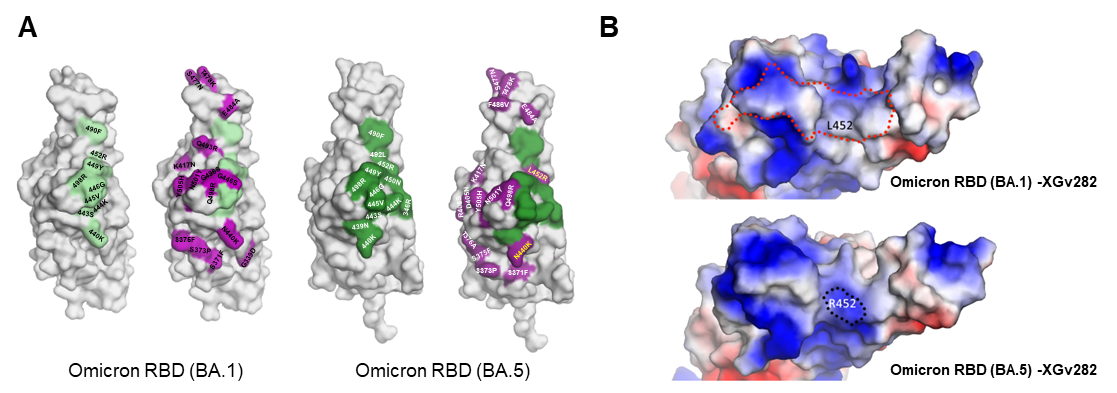

**Fig. S7**

Structural comparison between Omicron BA.1 RBD and Omicron BA.5 RBD in complex with XGv282.

**(A)** The epitopes of XGv282. These epitopes are colored palegreen on Omicron BA.1 RBD or green on Omicron BA.5 RBD. The mutated residues of Omicron BA.1 or BA.5 are colored deep purple. **(B)** Electrostatics of Omicron BA.1 RBD is changed by the L452R mutation in Omicron BA.5.

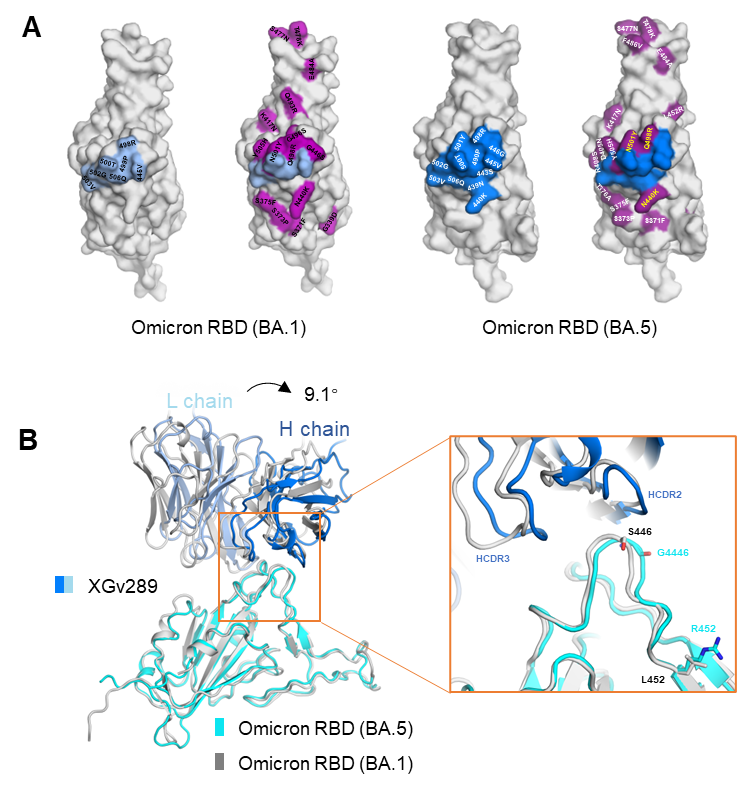

**Fig. S8**

Structural comparison between Omicron BA.1 RBD and Omicron RBD (BA.5) in complex with XGv289.

**(A)** The epitopes of XGv289. These epitopes are colored blue on Omicron BA.1 RBD or marine on Omicron BA.5 RBD. The mutated residues of Omicron BA.1 or BA.5 are colored deep purple. **(B)** Structural comparison of the Omicron BA.5 RBD-XGv289 complex and the Omicron BA.1 RBD-XGv289 complex (PDB ID: 7WE9). The RBD is superimposed. The boxed region is shown in inset in details.

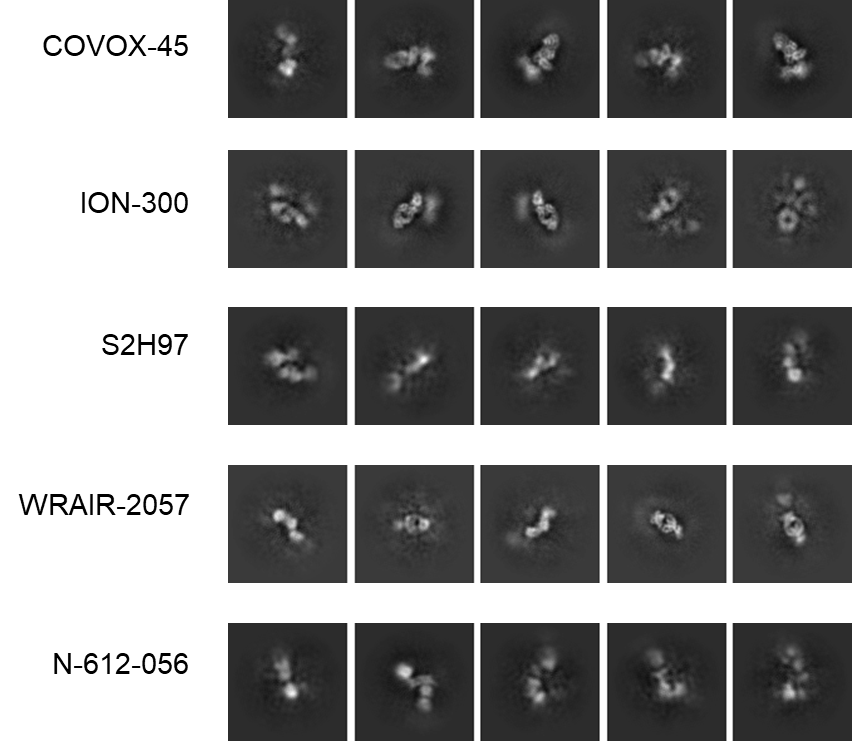

**Fig. S9**

Representative cryo-EM 2D class averages of Omicron BA.5 S in complex with antibodies of the Ab-H-RBD class.

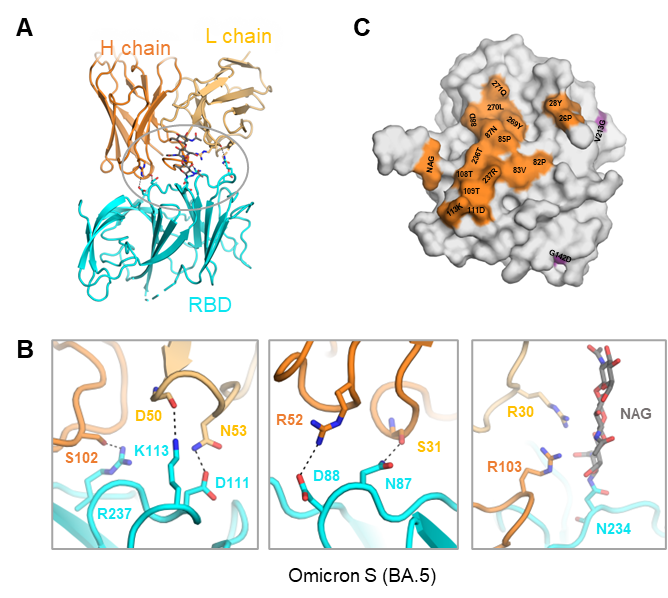

**Fig. S10**

The interactions between Omicron BA.5 NTD and S2L20.

**(A)** Binding interface between Omicron BA.5 NTD and S2L20. (**B**) Extensive hydrophilic interactions on the interface. Polar interactions are indicated by black dashed lines. **(C)** The epitopes of S2L20 on Omicron BA.5 RBD. These epitopes are colored orange. The mutated residues of Omicron BA.5 are colored deep purple.

**Table S1 | Data collection, 3D reconstruction and model statistics**

| **Data collection** |  |  |  |
| --- | --- | --- | --- |
| EM equipment | Titan Krios (Thermo Fisher Scientific) | | |
| Voltage (kV) | 300 | | |
| Detector | Gatan K3 Summit | | |
| Energy filter | Gatan GIF Quantum, 20 eV slit | | |
| Pixel size (Å) | 1.077 | | |
| Electron dose (e-/Å2) | 50 | | |
| Defocus range (μm) | -1.2 ~ -2.2 | | |
| Number of collected micrographs | 2,186 | 1,996 | 3,151 |
| Number of selected micrographs | 2,157 | 1,971 | 3,100 |
| Sample | S (BA.5) -XGv282 | S (BA.5) -XGv289 | S (BA.5) -S2L20 |
| PDB ID | 8GTO | 8GTP | 8GTQ |
| EMDB ID (whole map) | EMD-34259 | EMD-34261 | EMD-34263 |
| EMDB ID (local map) | EMD-34260 | EMD-34262 | EMD-34264 |
| **3D Reconstruction** |  |  |  |
| Software | cryoSPARC | Relion/cryoSPARC | cryoSPARC |
| Number of used particles | 350,760 | 137,483 | 162,448 |
| Resolution (Å) | 3.2 | 3.1 | 3.1 |
| Symmetry | C1 | | |
| Map sharpening B factor (Å^2^) | -90 | | |
| **Refinement** |  |  |  |
| Software | Phenix | | |
| Cell dimensions (Å) | 344.640 | 310.176 | 344.640 |
| Model composition |  |  |  |
| Protein residues | 3,762 | 3,768 | 3,759 |
| Side chains assigned | 3,762 | 3,768 | 3,759 |
| Sugar | 69 | 69 | 75 |
| R.m.s deviations |  |  |  |
| Bonds length (Å) | 0.006 | 0.005 | 0.005 |
| Bonds Angle (˚) | 1.036 | 0.853 | 0.862 |
| Ramachandran plot statistics (%) |  |  |  |
| Preferred | 92.29 | 94.17 | 95.14 |
| Allowed | 7.51 | 5.55 | 4.77 |
| Outlier | 0.20 | 0.28 | 0.10 |
